# Deletion of the 5-HT3A receptor reduces behavioral persistence and enhances flexibility

**DOI:** 10.1101/2025.09.02.673704

**Authors:** Tomoaki Nakazono, Makoto Kondo

**Author notes:** **Corresponding authors:** Tomoaki Nakazono and Makoto Kondo., **Email address:** (TN), (MK). Advanced Neuroimaging Center, National Institutes for Quantum Science and Technology, Chiba 263-8555, Japan (Tomoaki Nakazono).

## Abstract

The 5-HT3A receptor is the only ionotropic serotonin receptor and has been implicated in cognitive functions, yet its specific role remains unclear. To investigate the contribution of the 5-HT3A subtype, we trained wild-type (C57BL/6J) and 5-HT3A receptor knockout (Htr3a^−/−^) mice across a series of operant conditioning tasks and compared their behavioral performance. Following nose-poke training, both groups underwent a rule-switching task, extinction tests under fixed ratio (FR) and variable ratio (VR) reinforcement schedules, and a progressive ratio (PR) task to assess persistence. We found that 5-HT3A receptor knockout mice exhibited reduced responding during the extinction and PR tasks, suggesting diminished behavioral persistence. Notably, however, knockout mice acquired the new rule in the switching task significantly faster than wild-type controls, indicating enhanced cognitive flexibility. These findings suggest that the 5-HT3A receptor plays a role in regulating the balance between behavioral persistence and flexibility, normally biasing this balance toward persistence under normal physiological conditions. This mechanism may underlie the therapeutic effect of 5-HT3A receptor antagonists in treatment-resistant obsessive-compulsive disorder (OCD).

**Highlights:**

- Cognitive abilities of 5-HT3A receptor-deficient (Htr3a−/−) mice were examined using an operant conditioning paradigm.
- 5-HT3A receptor knockout mice exhibited reduced responding during extinction and progressive ratio (PR) tasks, indicating diminished behavioral persistence.
- 5-HT3A receptor knockout mice showed more rapid acquisition in a rule-switching task compared to wild-type mice, suggesting enhanced flexibility.
- The 5-HT3A receptor contributes to the balance between behavioral persistence and flexibility.
- This function may underlie the therapeutic effects of 5-HT3A receptor antagonists in treatment-resistant obsessive-compulsive disorder (OCD).

## 1. Introduction

Among the seven known subtypes of serotonin receptors, only the 5-HT3A receptor functions as a ligand-gated ion channel. Owing to this unique property, the 5-HT3A receptor is believed to mediate rapid synaptic transmission and contribute to rapid information processing in the brain [1]. In contrast, other serotonin receptors, which are G protein-coupled, primarily modulate slower, volume-based transmission and are more commonly associated with emotional regulation. Given its rapid signaling properties, the 5-HT3A receptor has been hypothesized to play a role in cognitive functions. Indeed, it is abundantly expressed in brain regions implicated in cognition, such as the prefrontal cortex and hippocampus [2]. However, the precise roles of 5-HT3A receptors in cognitive information processing remain largely unknown.

Several previous studies have reported potential roles of the 5-HT3A receptor in cognitive functions. In the context of fear conditioning, the 5-HT3A receptor is thought to be involved in the processing of fear-related memories [3–5], although a consistent mechanistic understanding has yet to be established. With respect to hippocampus-dependent spatial information processing, Yu et al. (2022)[6] demonstrated that 5-HT3A receptor knockout mice exhibited impaired spatial memory in the Morris water maze, while their object recognition performance remained intact.

Additionally, Smit-Rigter et al. (2010)[7] reported that the 5-HT3A receptor contributes to the regulation of social behavior.

Operant conditioning is considered a useful experimental paradigm for investigating the neuronal mechanisms underlying learning and memory. However, there have been relatively few studies focusing on the role of the 5-HT3A receptor in operant learning. Several earlier studies using non-human primates have reported that antagonists of the 5-HT3A receptor facilitate learning[8– 10]. For example, Barnes et al. (1990)[10] reported that ondansetron, a 5-HT3A receptor antagonist, enhanced reversal learning in marmosets. In contrast, Frick et al. (2015)[11] showed that ondansetron impaired acquisition and accelerated extinction in an operant task. More recently, Yoshida et al. (2019)[12] demonstrated that the 5-HT3A receptor in the rat ventral hippocampus plays an important role in sustaining goal-directed behavior.

Sustained goal-directed behavior is generally considered adaptive and essential for survival. However, when such persistence becomes excessive or inflexible, it can lead to maladaptive outcomes, as observed in psychiatric conditions such as obsessive-compulsive disorder (OCD). The practice guideline of the American Psychiatric Association (Koran et al., 2007)[13] even lists ondansetron as a potential augmentation strategy when selective serotonin reuptake inhibitors (SSRIs) are ineffective. Nevertheless, the precise role of the 5-HT3A receptor in the pathophysiology of OCD remains poorly understood.

To examine the role of the 5-HT3A receptor in both cognitive flexibility and persistence—traits potentially linked to OCD—we conducted a series of operant tasks in the same cohort of mice, enabling a multifaceted behavioral assessment. For this purpose, we employed the home-cage-compatible, standalone operant experimentation device FED3 [14,15]. Using this system, we performed reversal learning, extinction tests under two reinforcement schedules, and a progressive ratio schedule task [16]. We then compared learning ability, behavioral flexibility, and persistence between wild-type and 5-HT3A receptor knockout mice.

## 2. Materials and Methods

### 2.1.Animals

Twelve male 5-HT3A receptor knockout mice (5-HT3AR<sup>-/-</sup>)[17,18] were used as the experimental group. These mice were obtained from the Jackson Laboratory (Bar Harbor, ME, USA; strain #005251) and had been backcrossed to a C57BL/6J background for more than ten generations[19]. As controls, twelve male C57BL/6J mice (Japan SLC, Japan) were used. Only male mice were included in this study because previous reports have described sex differences in operant conditioning behavior [20]. Body weight and general health were carefully monitored throughout the experimental period. Experimental sessions were discontinued if an animal failed to consume enough pellets to maintain its body weight above 80% of its free-feeding weight. One wild-type mouse was excluded from the analysis for this reason. Mice were 15 to 20 weeks old at the beginning of behavioral training. Before the experiments, animals were group-housed with littermates; during the experimental period, they were housed individually. All mice were maintained under a 12 h light / 12 h dark cycle (lights on at 8:00 a.m.). All procedures were conducted in accordance with institutional and national guidelines and were approved by the Institutional Animal Care and Use Committee of Osaka Metropolitan University. Researchers were not blinded to group allocation.

### 2.2 Apparatus

Each mouse was provided with two cages (Optimice; Animal Care Systems, Centennial, CO, USA): a home cage and an experimental cage. The home cage contained paper pulp chip bedding (ALPHA-dri; Shepherd Specialty Papers, USA) and no additional objects. The experimental cage was of the same type as the home cage. Inside the experimental cage, we installed a standalone, wireless operant conditioning system (FED3)[14,15]. A custom-made acrylic partition was placed to prevent the animal from accessing the rear of the device. Sheet bedding (ALPHA Pad; Shepherd Specialty Papers, USA) was used instead of chip bedding to prevent pulp chips from entering the pellet dispenser. Animals had ad libitum access to water in both cages. Both cages were placed in the same animal room.

The FED3 device features two nose-poke ports equipped with infrared sensors and a central pellet magazine that delivers 20 mg precision pellets as rewards. Behavioral tasks were controlled by custom Arduino programs. All behavioral events were logged to microSD cards and analyzed using custom scripts written in MATLAB (R2022a, MathWorks, Natick, MA) and R (version 4.3.1; R Core Team, 2023).

### 2.3. Training and Behavioral Tasks

Before training, animals were group-housed with free access to food. One day prior to the start of training, animals were singly housed in the home cage, and their body weight was recorded to define the free-feeding weight. On the same night, 20 mg precision pellets were provided in the home cage to allow habituation to the novel food. Behavioral sessions were conducted daily between 10:00 and 18:00. At the beginning of each session, animals were transferred from the home cage to the experimental cage and returned afterward. Body weight was recorded before and after each session.

The timeline of the experimental procedures is illustrated in Figure 1, as described in the following sections.

**Figure 1.**
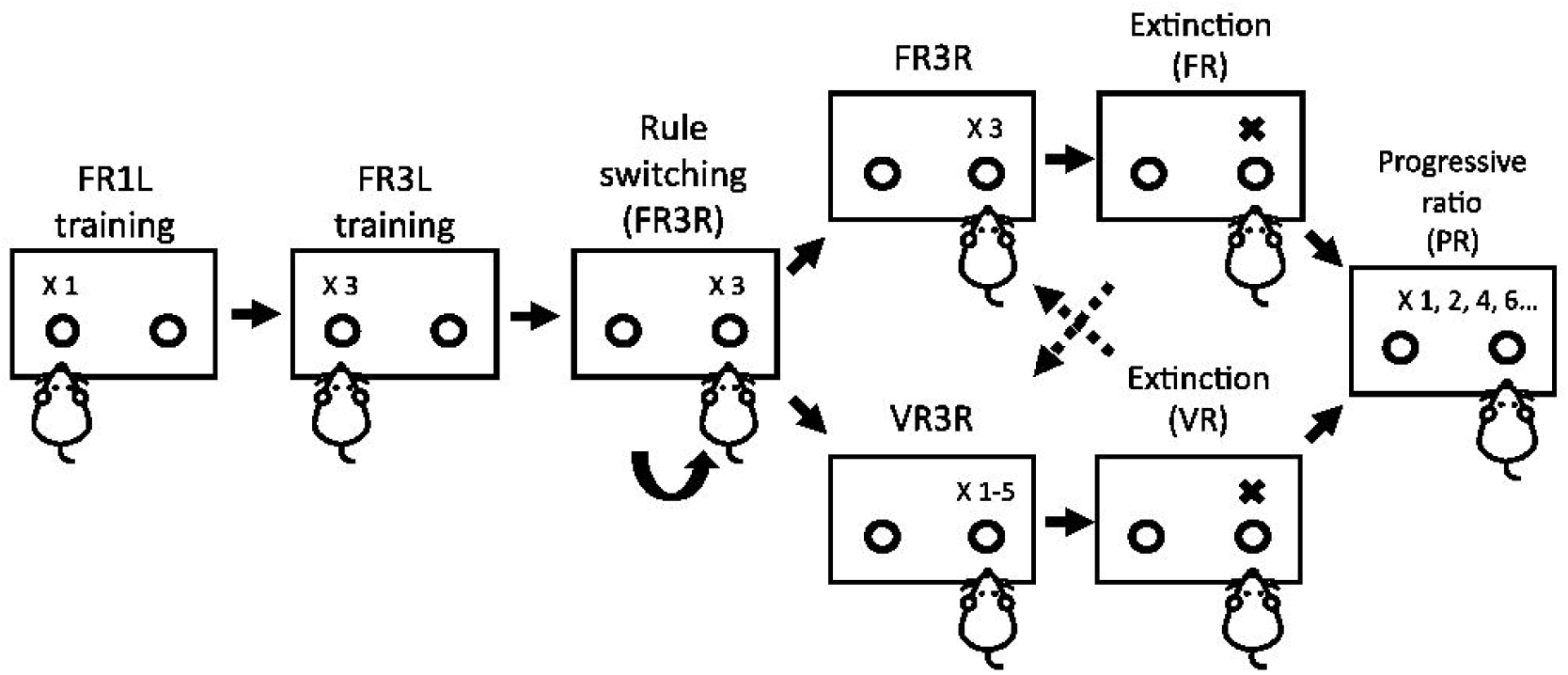
Overview of the experimental procedures. Mice were trained and tested across multiple operant conditioning tasks using a standalone behavioral apparatus equipped with two nose-poke ports (FED3.1).

#### 2.3.1 Free Feeding Training

In this phase, the FED3 delivered a single precision pellet to the magazine. When the animal retrieved the pellet (detected via the IR sensor), the next pellet was immediately delivered. Most animals learned to eat from the magazine within one day.

#### 2.3.2. Fixed Ratio Training (FR1L/FR3L)

Fixed ratio (FR) training began with an FR1 schedule, in which one nose-poke to the left port resulted in pellet delivery. Starting from this training phase, pellet rewards were accompanied by a 1-second tone and LED cue. When an animal obtained more than 100 pellets in a single session, FR3L training began the following day. Under the FR3 schedule, three left nose-pokes were required for pellet delivery. Nose-pokes to the right port were recorded but not reinforced. FR3L training continued until each animal reliably obtained sufficient pellets for three consecutive days.

#### 2.3.3 Rule-switching Task

In the rule-switching task, reinforcement was shifted to the right port using an FR3 schedule (FR3R), while left nose-pokes were no longer rewarded. No explicit cues signaled this rule switching. Accuracy was calculated as the ratio of nose-pokes into the correct port within a moving window of 30 consecutive nose-pokes. The learning criterion was defined as the first trial at which accuracy reached 80%.

#### 2.3.4. Extinction Test

Each animal underwent two extinction tests—Extinction 1 and Extinction 2—conducted under different reinforcement schedules: a fixed ratio (FR) schedule and a variable ratio (VR) schedule. The order of schedules was counterbalanced across subjects. Mice were divided into two groups: FR→VR (FV) and VR→FR (VF). In the FV group, animals received three days of FR3R training prior to Extinction 1, followed by three days of VR3R training prior to Extinction 2. The VR3R schedule delivered rewards after a random number of right nose-pokes, ranging from 1 to 5 (average = 3). In both extinction sessions, five pellets were delivered at the beginning of the session according to the previous schedule, followed by a 6-hour extinction period with no reinforcement. After Extinction 1, the FV group received VR training, and the VF group received FR training. After Extinction 2, all animals received FR training. To evaluate extinction speed, cumulative nose-poke responses during the first 120 minutes were plotted for each animal. The area under the curve (AUC) of these cumulative response plots was used as an index of extinction speed. AUC values were normalized to a maximum of 1 for comparison across animals.

#### 2.3.5. Progressive Ratio Test (PR Test)

The PR test was conducted to assess goal-directed motivation[21]. Following Extinction 2, all animals underwent one day of FR3R training before the PR test. In the PR test, reinforcement was provided at increasing response requirements, defined by the following equation:

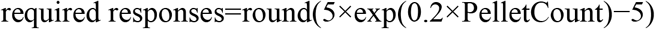

In this formula, the number of responses required for each subsequent reward based on the number of pellets already obtained (PelletCount). As a result, the response requirement increased in a nonlinear manner (e.g., 1, 2, 4, 6, 9, 12, 15, …), thereby imposing progressively greater effort on the animal. The PR session lasted 6 hours.

### 2.4. Statistical analysis

Statistical analyses were performed using R (version 4.3.1; R Core Team, 2023). For FR1 training data and extinction test data, repeated-measures analysis of variance (ANOVA) was conducted. For the comparison of learning speed in the rule-switching task, the Mann–Whitney U test was applied due to the presence of an extreme value in the WT group. For other comparisons between genotypes, unpaired t-tests with Welch’s correction were used. To examine the relationships between learning speed and multiple indices of extinction speed, we performed linear regression analyses. For correlation and regression analyses, data points that were deemed outliers based on extreme values in the switching task (e.g., learning speed > 600 trials) were excluded to avoid undue influence on model fitting and correlation coefficients. All results are presented as mean ± standard error of the mean (SEM). Statistical significance was defined as *p* < 0.05.

## 3. Results

### 3.1. Comparable body weight and food intake across genotypes

To rule out the possibility that differences in operant conditioning performance were due to motivational factors such as hunger level, we compared body weight and food intake on the day before behavioral testing. There were no significant differences between the two groups in body weight (*t* = −0.74, *df* = 21.00, *p* = 0.469; Fig. 2A) and food intake (*t* = 1.15, *df* = 15.76, *p* = 0.268; Fig. 2B), suggesting that both mouse groups were similarly motivated. We also compared body weights on each test day and found no significant differences (Supplementary Fig. 1).

**Figure 2.**
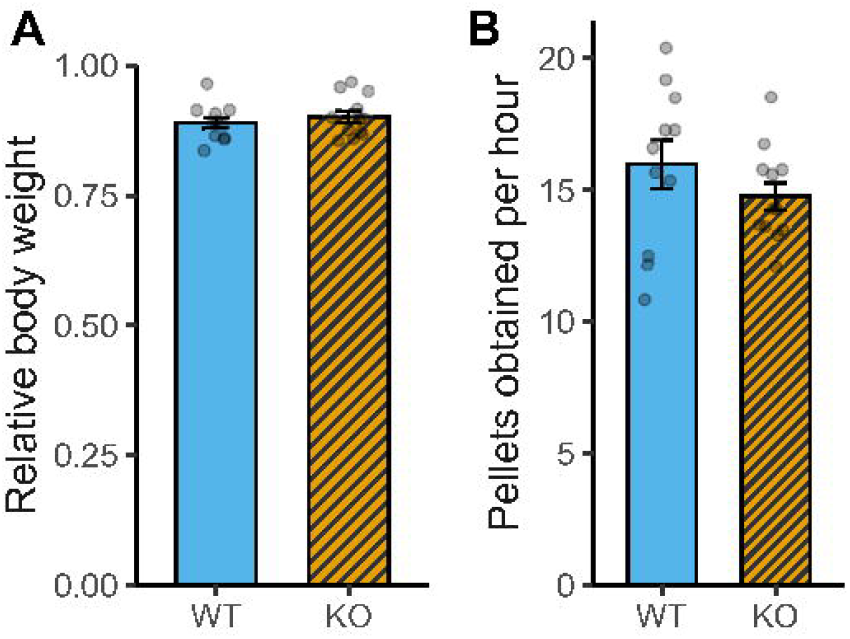
Body weight and food intake on the day before behavioral testing. (A) Mean body weight on the day before the switching task (i.e., the third day of FR3L training) relative to free-feeding weight. Body weight was measured immediately before the training session (*p* = 0.469). (B) Number of food pellets obtained per hour during the same training session (*p* = 0.268). Sample sizes: WT *n* = 11, KO *n* = 12.

### 3.2. 5-HT3A receptor knockout mice exhibit faster learning in the rule-switching task

Previous studies have reported that 5-HT3A receptor antagonists can affect performance in operant conditioning tasks, although the findings remain inconsistent.

As a baseline measure, we first examined whether there were any differences in learning speed during the initial fixed ratio (FR1) training phase. No significant differences were observed between the groups (Supplementary Fig. 2).

To assess the potential role of 5-HT3A receptors in cognitive flexibility, we then compared the number of trials required to reach 80% accuracy in a rule-switching task. Notably, KO mice exhibited a steeper learning curve (Fig. 3A, B) and reached the performance criterion with significantly fewer trials than WT mice (*U* = 111, *p* = 0.005; Fig. 3C). The effect was robust to exclusion of a single WT outlier.

**Figure 3.**
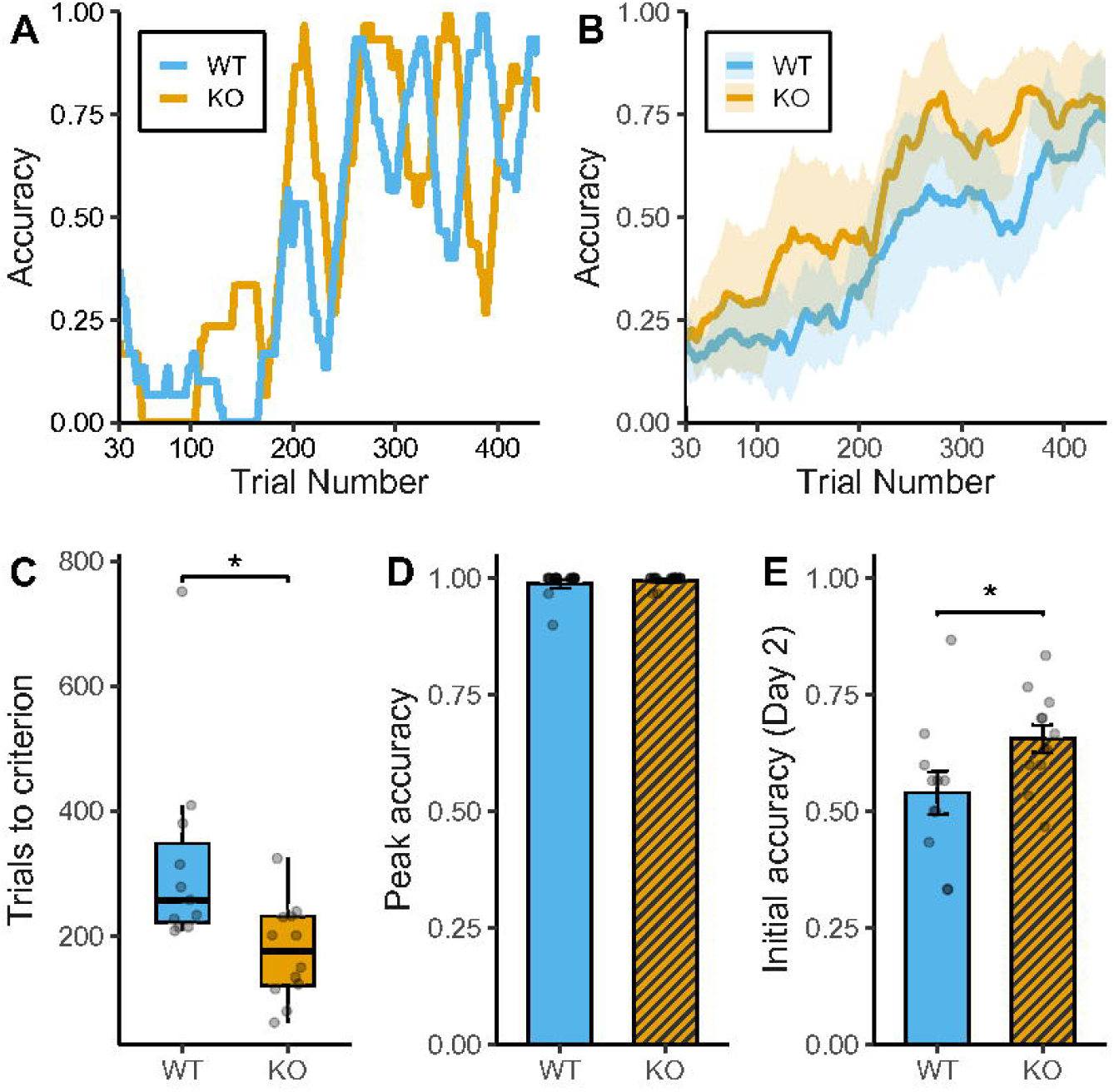
Performance in the rule-switching task. (A, B) Learning curves in the switching session. (A) Representative curves from one WT and one KO mouse. (B) Group mean curves with 95% confidence intervals. Blue and orange lines represent WT and KO mice, respectively. (C) Number of trials required to reach the 80% accuracy criterion. KO mice showed faster learning than WT mice (*p* = 0.005). Data are shown as box plots with individual data points overlaid. (D) Peak accuracy within the switching (*p* = 0.524). (E) Accuracy during the first 30 trials on the day following the rule switch. KO mice showed better performance than WT mice (*p* = 0.046). Sample sizes: WT *n* = 11, KO *n* = 12.

There was no significant difference in peak performance between the two groups, with both WT and KO mice achieving nearly 100% accuracy at their best (*t* = –0.66, *df* = 13.20, *p* = 0.524; Fig. 3D), indicating that both groups successfully acquired the task, albeit at different speeds.

Furthermore, we examined whether the effects of learning persisted into the following day by comparing accuracy during the initial 30 trials of the next session. KO mice performed significantly better than WT mice in this early phase (*t* = –2.15, *df* = 17.22, *p* = 0.046; Fig. 3E), indicating that differences in performance were also observed at the beginning of the subsequent session.

### 3.3. 5-HT3A receptor knockout mice show reduced persistence in extinction

To assess behavioral persistence, we employed two distinct paradigms: extinction testing and the progressive ratio (PR) schedule. While the extinction test evaluates the persistence of previously reinforced behavior in the absence of reward, the PR task assesses the subject’s willingness to sustain effortful responding under increasing response requirements. In the extinction test, two types of reinforcement schedules were examined, as resistance to extinction is well known to vary depending on the schedule, with variable ratio (VR) schedules typically producing greater resistance than fixed ratio (FR) schedules[22]. In the extinction test, a three-way repeated-measures ANOVA with Genotype (WT vs. KO) as a between-subjects factor and Schedule (FR vs. VR) and Time (1–6) as within-subjects factors revealed a significant main effect of Time, *F*(1.37, 28.86) = 65.19, *p* < 0.001. No significant main effects of Genotype, *F*(1, 21) = 0.43, *p* = 0.518, or Schedule, *F*(1, 21) = 0.21, *p* = 0.648, were observed, and no interactions were significant (Fig. 4A).

**Figure 4.**
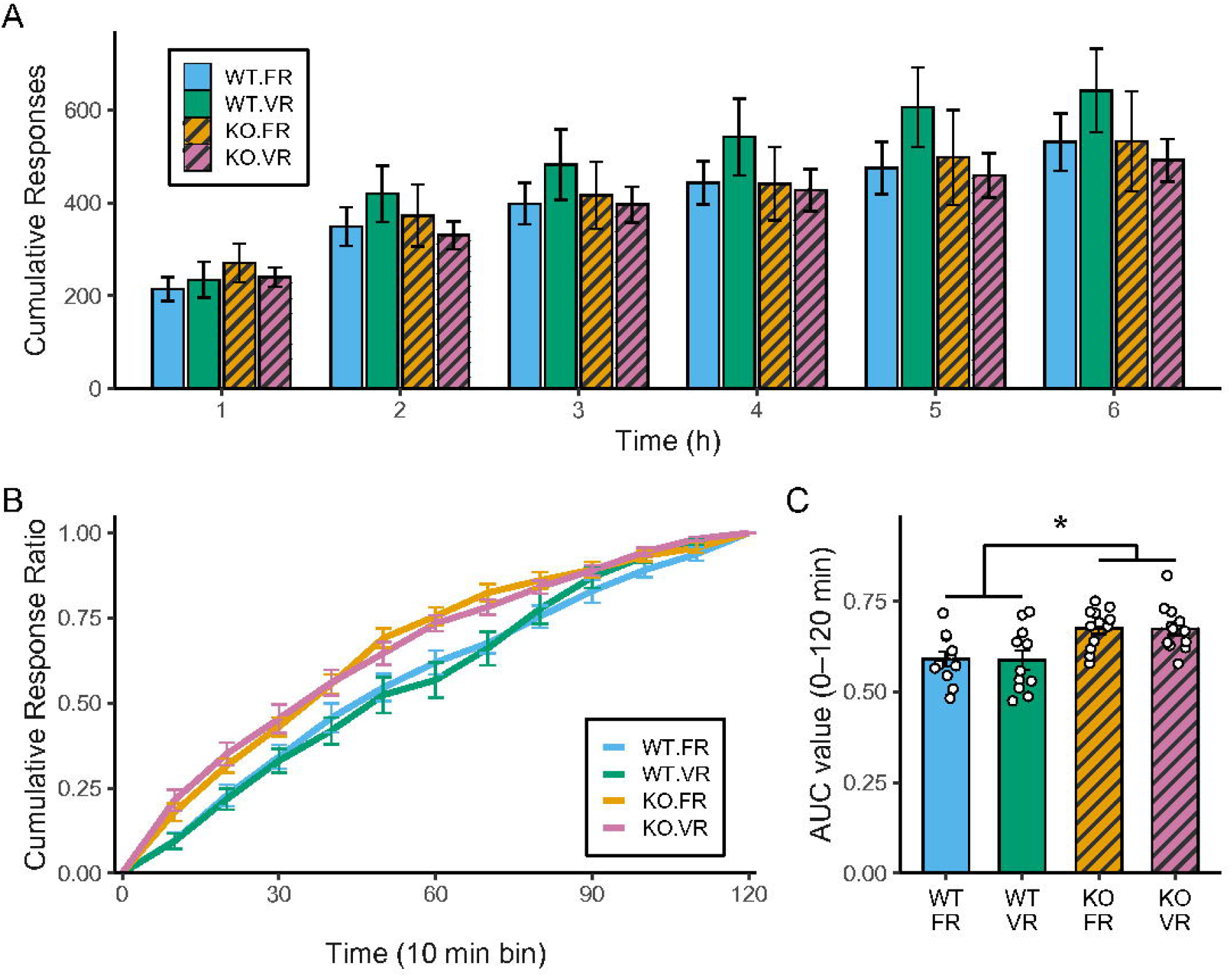
Genotype-dependent differences in extinction speed. (A) Cumulative responses per hour following the onset of extinction. A three-way repeated-measures ANOVA with Genotype as a between-subjects factor and Schedule and Time as within-subjects factors revealed a significant main effect of Time (*p* < 0.001). No significant main effects of Genotype (*p* = 0.518) or Schedule (*p* = 0.648) were observed, and no interactions were significant. (B) Time course of the cumulative response ratio during the first 120 minutes of the extinction session. (C) Comparison of the area under the curve (AUC) value for cumulative response ratios over the first 120 minutes. A two-way repeated-measures ANOVA with Genotype as a between-subjects factor and Schedule as within-subjects factors revealed a significant main effect of Genotype (*p* < 0.001). No significant main effect of schedule was observed (*p* = 0.916) and interaction was no significant (*p* = 0.981). Sample sizes: WT *n* = 11, KO *n* = 12.

To evaluate the speed of extinction, we analyzed the cumulative response ratio over the first 120 minutes of the session, as responding decreased markedly beyond this point. KO mice showed a trend toward faster extinction compared with WT mice, as evidenced by the trajectory of cumulative response ratios (Fig. 4B). The area under the curve (AUC) for this period was significantly greater in KO mice, indicating a faster decline in responding (Genotype: *F*(1, 21) = 16.83, *p* < 0.001; Fig. 4C). No significant effect of schedule was observed (*F*(1, 21) = 0.01, *p* = 0.916).

In the PR test, we compared the mean number of pellets obtained during the session. Although KO mice tended to obtain fewer pellets, the difference did not reach statistical significance (*t* = 1.90, *df* = 20.80, *p* = 0.071; Fig. 5).

**Figure 5.**
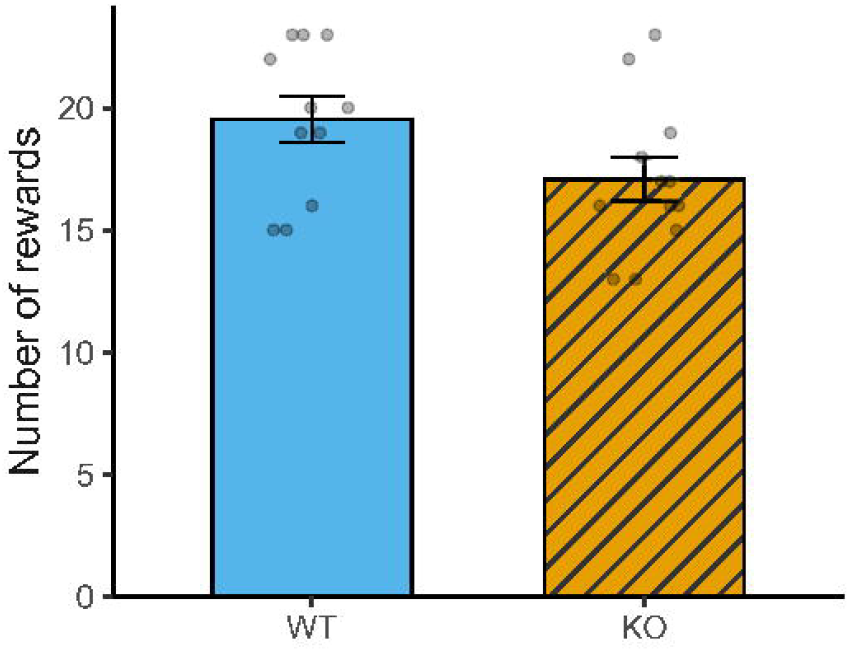
Genotype differences in reward acquisition in the progressive ratio task. Each bar represents the mean total number of pellets obtained at the 6-hour time point in the progressive ratio task (± SEM). Individual data points are shown. No significant difference was observed between WT and KO mice (*p* = 0.07). Sample sizes: WT *n* = 11, KO *n* = 12.

### 3.4. No relationship between learning speed and extinction

Finally, we examined whether learning speed in the rule-switching task was associated with extinction-related measures (Fig. 6). The number of trials required to reach the learning criterion was used as an index of learning speed. The AUC values under both FR and VR schedules, as well as the number of rewards obtained in the PR task, were used as indices of extinction speed or persistence. Regression analyses revealed no significant relationships between learning speed and any of these measures.

**Figure 6.**
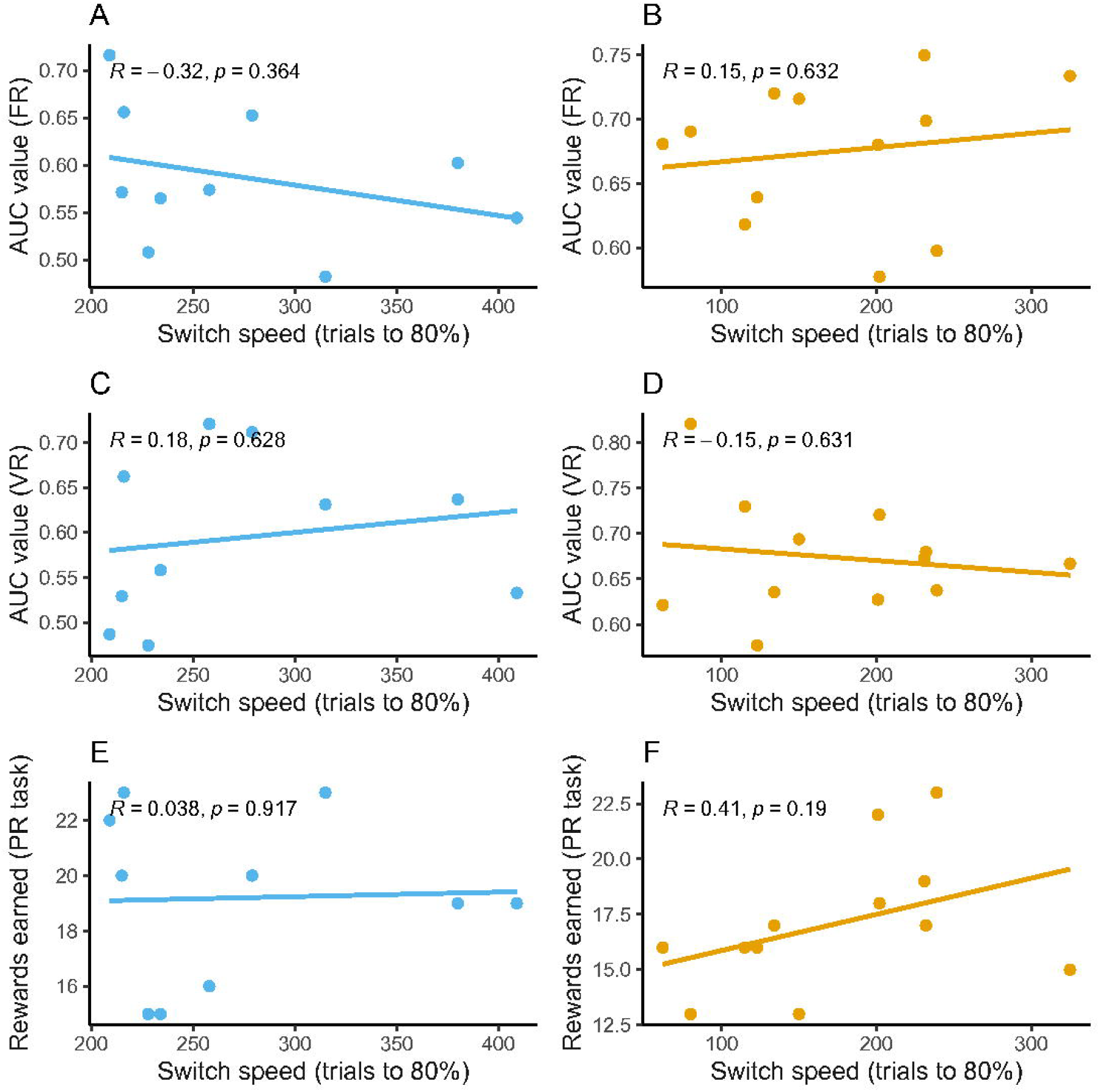
No clear relationships between learning speed in the rule-switching task and extinction or motivation measures. Scatter plots show the relationship between learning speed and the area under the extinction curve (AUC) values under the FR schedule (top row), the AUC values under the VR schedule (middle row), and the number of rewards earned in the progressive ratio (PR) task (bottom row). The left column shows data from WT mice, and the right column shows data from KO mice. Each dot represents an individual mouse. Solid lines indicate linear regression fits, with the results of regression analyses shown in the upper left corner of each plot. Data points that were deemed outliers based on extreme values in the switching task were excluded to avoid undue influence on model fitting and correlation coefficients. Sample sizes: WT *n* = 10, KO *n* = 12.

## 4. Discussion

In this study, we investigated the role of the 5-HT3A receptor in operant conditioning by comparing WT mice and KO mice. There were no significant genotype differences in food intake or motivational levels (Fig. 2). Although the 5-HT3A receptor is also expressed in the gastrointestinal system and its antagonists are widely used as antiemetics, KO mice exhibited comparable feeding-related motivation in operant tasks. In the rule-switching task, KO mice successfully acquired the new rule and showed faster learning than WT mice (Fig. 3), suggesting enhanced behavioral flexibility. In contrast, KO mice exhibited faster extinction (Fig. 4) and obtained fewer rewards in the progressive ratio task (Fig. 5), suggesting reduced behavioral persistence. At the individual level, no significant correlation was observed between behavioral flexibility and persistence (Fig. 6).

Serotonin is known to play a key role in regulating motivation. However, the effects of serotonin on motivation differ across its receptor subtypes. For example, Hori et al. (2024)[23] demonstrated that both 5-HT1A and 5-HT1B receptors contribute to motivational processes, but in distinct ways. In contrast, our results suggest that the 5-HT3A receptor may not be directly involved in feeding-related motivation itself (Fig. 2).

Serotonin is also considered to be important for behavioral persistence [24,25]. Yoshida et al. (2019)[12] reported that 5-HT3A receptors expressed in ventral hippocampal interneurons play a critical role in sustaining persistent behavior. Consistent with this, our extinction data also suggests a similar effect: the faster extinction observed in 5-HT3A receptor KO mice (Fig. 4) may reflect a deficit in the ventral hippocampal function. In addition, Miyazaki et al. (2020)[26] demonstrated that serotonergic projections from the dorsal raphe nucleus (DRN) to the orbitofrontal cortex contribute to waiting for delayed rewards. It is therefore possible that 5-HT3A receptors are also involved in this pathway and influence persistence through prefrontal mechanisms.

In fear conditioning (Pavlovian conditioning), a previous study reported that 5-HT3A receptor KO mice showed slower extinction than wild-type mice[5]. Taken together, the Pavlovian finding and our operant results suggest that that 5-HT3A receptor has opposite roles in Pavlovian conditioning and in operant conditioning because different neuronal circuits are involved in Pavlovian and operant conditioning[27].

We did not detect a significant effect of the reinforcement schedule on extinction speed (Fig. 4A). However, WT mice tended to emit more responses over the 6-h test than other conditions in VR schedule. It suggested that WT mice in VR schedule had a tendency to continue to nose-poking through the session. This tendency is consistent with the theory that VR schedules confer greater resistance to extinction[22] whereas KO mice did not exhibit this tendency. Miyazaki et al. (2020)[26] suggested that different neural pathways are engaged depending on the certainty of the expected reward. They reported that serotonergic projections from the dorsal raphe nucleus (DRN) to the orbitofrontal cortex (OFC) are preferentially involved under high-certainty conditions, whereas projections to the medial prefrontal cortex (mPFC) are engaged under conditions of reward uncertainty. Given that the FR schedule can be considered a high-certainty condition and the VR schedule an uncertain one, it is possible that the role of the 5-HT3A receptor is also modulated by reward certainty.

Persistence is often regarded as a key to success, and its positive aspects are typically emphasized. However, excessive persistence can be maladaptive in certain contexts and has been implicated in psychiatric conditions such as OCD. Our findings from the rule-switching task and extinction test suggest a trade-off between behavioral persistence and cognitive flexibility. While 5-HT3A receptor KO mice exhibited reduced persistence, they adapted to new rules more rapidly than WT mice (Fig. 3). These results suggest that the 5-HT3A receptor may play a key role in regulating the balance between persistence and flexibility.

Interestingly, several studies have suggested that serotonin is involved in behavioral flexibility. Clarke et al. (2004)[28] reported that serotonin depletion in the prefrontal cortex of marmosets increased behavioral persistence. Previous studies also demonstrated that serotonergic projections from the DRN to the OFC facilitated flexible reversal learning [29,30]. In contrast, a depletion study in rats showed increased win–stay behavior and decreased lose–shift behavior following serotonergic disruption [31]. A pharmacological study using a 5-HT3A receptor antagonist also reported findings consistent with our results [10]. These mixed findings support the hypothesis that serotonin contributes to the flexibility–persistence trade-off, but its effects may differ depending on the receptor subtype. While most serotonin receptors appear to promote flexibility, the 5-HT3A receptor may play an opposing role that increases persistence and reduces flexibility. Unlike other serotonin receptors, the 5-HT3A receptor is a ligand-gated ion channel rather than a G-protein-coupled receptor, which may underlie its distinct contribution to neural computation and behavior.

This hypothesis is consistent with the fact that 5-HT3A receptor antagonists are considered potential therapeutic options for patients with OCD who do not respond to SSRIs [13]. In fact, several meta-analyses have demonstrated the efficacy of 5-HT3A receptor antagonists in treating OCD [32,33]. However, little is known about the mechanisms by which 5-HT3A receptors contribute to the pathophysiology of OCD. Hatakama et al. (2022)[34] suggested that the treatment effects of SSRIs in OCD occur through the 5-HT2C receptor. Our results suggest that serotonin may affect OCD via multiple pathways and receptor subtypes. Understanding the cognitive roles of 5-HT3A receptor antagonists may help in developing novel therapeutic strategies for treatment-resistant OCD.

Interaction with the dopaminergic system is a possible mechanism underlying the function of the 5-HT3A receptor. Many studies have shown that there are strong interactions between the serotonergic and dopaminergic systems [35]. Cinotti et al. (2019)[36] demonstrated that dopamine blockade in rats induced random choice behavior and proposed that dopamine plays a key role in regulating the exploration–exploitation trade-off. More recently, Ishino et al. (2023)[37] reported a novel property of dopamine neurons that promotes retry behavior following unrewarded outcomes. These findings raise the possibility that serotonergic pathways may interact with such dopaminergic circuits, and that the 5-HT3A receptor may play a role in modulating this interaction.

It also remains an open question whether the bidirectional modulation between flexibility and persistence is mediated by the same neuronal pathways. Kawai et al. (2022)[38] reported that projections from the median raphe nucleus (MRN) contribute to negative information processing, whereas serotonergic pathways from the dorsal raphe nucleus (DRN) are involved in processing positive information. It is also possible that the DRN and MRN contribute differentially to behavioral regulation along the flexibility–persistence axis. If flexibility and consistency were processed by different serotonergic pathways, it might explain the lack of correlation between performance in the flexibility and persistence tasks in this study (Fig.6).

In this study, we used a standalone operant conditioning system [14,15]. This type of setup allowed for extended behavioral sessions in a context similar to the home cage. Under mild food restriction, mice were able to perform more than 100 trials per session (Fig. 2). Such mild restriction is not only preferable from an animal welfare perspective, but also improves experimental efficiency, as usable data were obtained from 23 out of 24 mice. This system is well suited for conducting multiple cognitive tasks, enabling the exploration of relationships between different cognitive domains (Fig.1).

However, one limitation of this approach is the potential for task interference, as sequential testing may lead to carry-over effects. In our study, we did not observe a significant genotype difference in the PR task (Fig. 5), which may have been influenced by the two preceding extinction sessions. Another limitation of our study is that, because we used global knockout animals, we were unable to identify the specific brain regions or neural circuits in which the 5-HT3A receptor is involved. To address these issues, future studies should combine operant tasks with other techniques, such as optogenetic manipulation or region-specific pharmacological interventions.

In conclusion, this study demonstrated that the 5-HT3A receptor plays a role in regulating the balance between behavioral persistence and flexibility. Our findings suggest that the 5-HT3A receptor biases behavior toward persistence. Future work using targeted genetic or circuit-level manipulations will help elucidate the neural basis of 5-HT3A receptor function and its relevance to the pathophysiology of treatment-resistant obsessive-compulsive disorder.

## Acknowledgements

We thank all members of our laboratory for their insightful comments and valuable assistance with animal care.

## Declaration of generative AI and AI-assisted technologies in the writing process

During the preparation of this manuscript, the authors used ChatGPT to assist with language editing. After using the tool, the authors carefully reviewed and revised the content as necessary and take full responsibility for the final version of the manuscript.

## Data availability Statement

Data will be made available on request.

## CRediT author statement

Tomoaki Nakazono: Conceptualization, Software, Formal analysis, Investigation, Writing - Original Draft, Writing - Review & Editing, Visualization, Project administration, Funding acquisition. Makoto Kondo: Conceptualization, Resources, Writing - Review & Editing, Supervision, Project administration, Funding acquisition.

## Figure captions

**Supplementary Figure 1.**
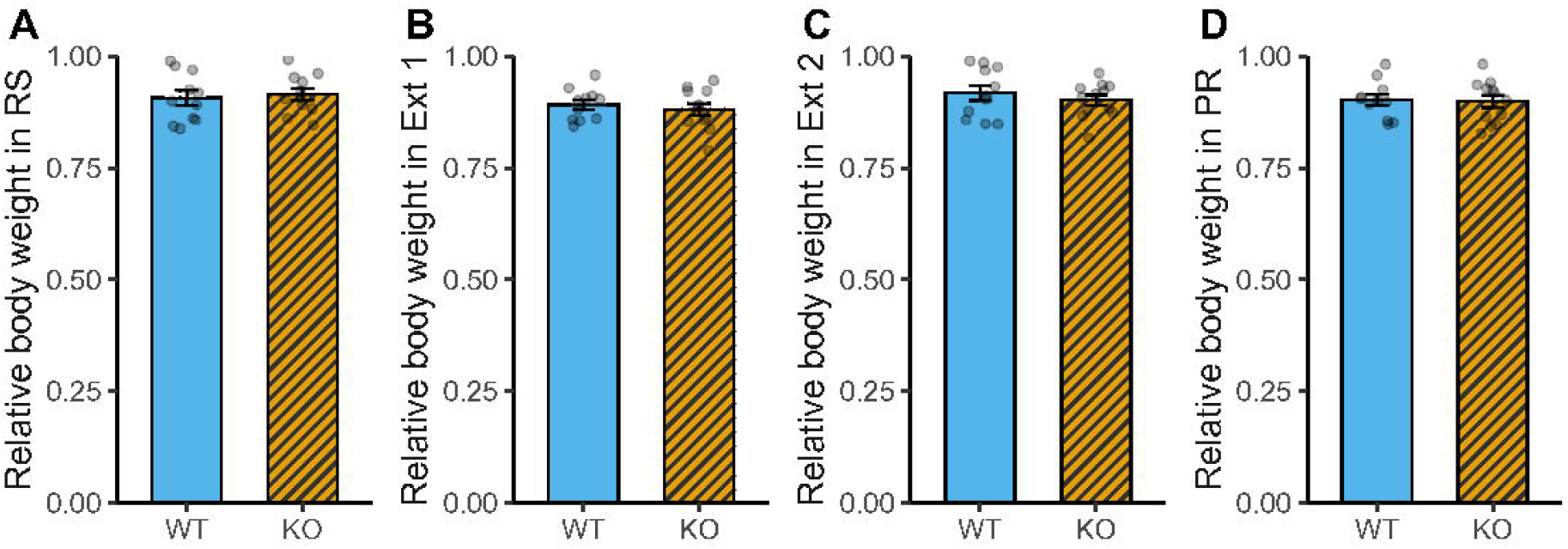
Body weights on the days of behavioral tests. Mean body weights on the days of each behavioral task, expressed relative to free-feeding weight. (A) Rule switching test (*p* = 0.469), (B) Extinction test 1 (*p* = 0.544), (C) Extinction test 2 (*p* = 0.442), (D) Progressive ratio task (*p* = 0.850). Sample sizes: WT *n* = 11, KO *n* = 12.

**Supplementary Figure 2.**
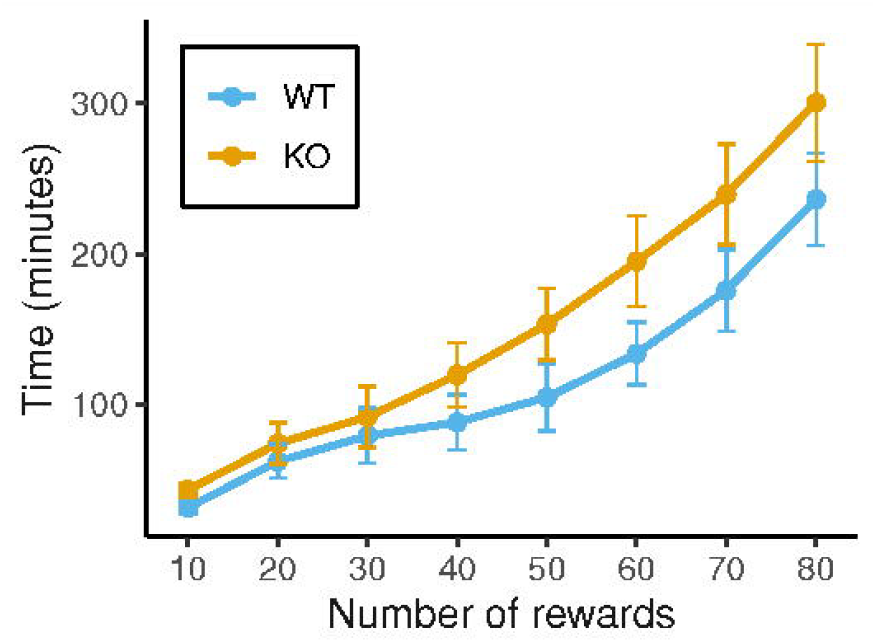
Learning performance during initial FR1 training. The graph shows the cumulative time (in minutes) required to obtain rewards, plotted against the number of rewards earned in increments of 10. Data are presented as group means. Blue and orange lines indicate WT and KO mice, respectively. A two-way ANOVA revealed no significant main effect of Genotype on the time required to obtain rewards during FR1 training (*F*(1, 21) = 1.15, *p* = 0.296). Sample sizes: WT *n* = 11, KO *n* = 12.

